# Ca²⁺ oscillations promote microtubule-band turnover and support tip growth in Arabidopsis zygotes

**DOI:** 10.64898/2026.02.19.706723

**Authors:** Hikari Matsumoto, Zichen Kang, Tomonobu Nonoyama, Yusuke Kimata, Satoru Tsugawa, Minako Ueda

## Abstract

The zygote is the origin of development, and in most angiosperms, it divides asymmetrically to establish the apical–basal axis. In *Arabidopsis thaliana*, the zygote undergoes tip growth-like polar elongation, using a subapical transverse microtubule band (MT band). Because canonical tip-growing cells rely on longitudinal actin filaments (F-actin), it remains unclear whether and how the zygote employs conserved tip-growth mechanisms. Here, using quantitative live-cell imaging, pharmacological perturbations, and mechanical simulations, we found that oscillatory Ca²⁺ waves, a hallmark of tip growth, are coupled to zygote elongation through a bidirectional feedback loop, as in other tip-growing cells. However, Ca²⁺ waves were dispensable for overall F-actin alignment but promoted MT band turnover. Our study provides a model showing that the zygote uses a conserved tip-growth module, in which Ca²⁺ oscillations and cell elongation reinforce each other, but redirects the target cytoskeleton to the MT band, enabling zygote-specific tip growth for axis formation.

## Introduction

Axis formation is an essential event in the development of multicellular organisms. In angiosperms, the apical–basal (shoot–root) axis is generally established by the polarity of the zygote. In *Arabidopsis thaliana* (Arabidopsis), the zygote undergoes polar cell elongation toward the apical side and then divides asymmetrically to generate a small apical cell and a large basal cell ^1^. The apical cell gives rise to most of the plant body, whereas the basal cell produces part of the root tip and the extraembryonic suspensor.

Recent live-cell imaging and image-analysis approaches have revealed that the zygote elongates at the apical end, where the sperm cell fused to the egg cell, in a manner of tip growth, a mode in which growth is confined to the apical tip ^2^. In canonical tip-growing cells, such as pollen tubes and root hairs, cytosolic calcium (Ca²⁺) concentrations at the growing apex repeatedly increase and decrease, forming Ca²⁺ oscillations ^3,4^, and Ca²⁺ signals regulate alignment of actin filaments (F-actin) that support sustained polar cell elongation ^5,6^. In the zygote, F-actin aligns along the apical–basal axis, similar to the longitudinal array along the growth direction in typical tip-growing cells. However, in the zygote, treatment with actin polymerization inhibitors has little effect on cell elongation, while it severely blocks polar organelle migration in zygotes ^7,8^.

In contrast, microtubule (MT) organization is directly linked to zygote elongation through mechanical feedback; the zygote possesses a transverse band of cortical MTs in the subapical region, termed the MT band, which forms a mechanical frame that restricts lateral cell expansion and thereby drives polar elongation toward the tip ^9^. Zygote elongation, in turn, increases estimated surface tension that guides MT assembly, enabling the MT band to be maintained at an appropriate width and at a constant distance from the tip during zygote elongation ^9,10^. These MT dynamics are apparently different in pollen tubes, in which MTs align along the growth direction, similar to the longitudinal F-actin^11^. This difference in cytoskeletal features raises the question of whether the zygote uses the canonical tip-growth Ca²⁺ module and, if so, how it is redirected to support MT-dependent tip growth in the zygote.

Here, by combining high-resolution live-cell imaging of multicolor fluorescent markers, quantitative image analyses, pharmacological perturbations, and mechanical simulations, we discovered that oscillatory Ca²⁺ waves occur in the Arabidopsis zygote. We further show that Ca²⁺ oscillations are tightly coupled to, and required for zygote elongation, indicating a bidirectional feedback loop. In addition, we show that Ca²⁺ oscillations are dispensable for the longitudinal F-actin alignment but contribute to MT band turnover and thereby to proper maintenance of the MT band during zygotic tip growth.

## Results

### Oscillatory Ca²⁺ waves arise at the apical tip of the zygote

To examine whether Ca²⁺ oscillations occur in the zygote, we expressed the MatryoshCaMP6s reporter ^12^, which combines a green fluorescent Ca²⁺ sensor (GCaMP6s) with an orange cytosolic reference, under the egg cell- and zygote-specific *EGG CELL1* (*EC1*) promoter (EC1p::MatryoshCaMP6s). We initially combined a red plasma membrane (PM) marker (EC1p::tdTomato-SYP121) to generate Ca²⁺/cytosol (cyto)/PM marker line. Using two-photon excitation microscopy (2PEM), we detected Ca²⁺ signal in the apical region of the zygote (Fig. 1a). However, because the cytosolic reference and the PM marker were not spectrally separable (Cyto+PM in Fig. 1a), we instead used a cyan nuclear marker (EC1p::H2B-mTFP) to generate a Ca²⁺/cyto/nucleus marker line. In this line, the bright cyan fluorescence from the nuclear marker bled into the Ca²⁺ detection channel, resulting in an apparent nuclear signal (Ca²⁺(+nuc) in Fig. 1b). Despite this, apical Ca²⁺ fluorescence was consistently detected, and the nuclear marker was useful for staging zygotes (see below). Therefore, we primarily used the Ca²⁺/cyto/nucleus marker line for subsequent analyses.

**Figure 1.**
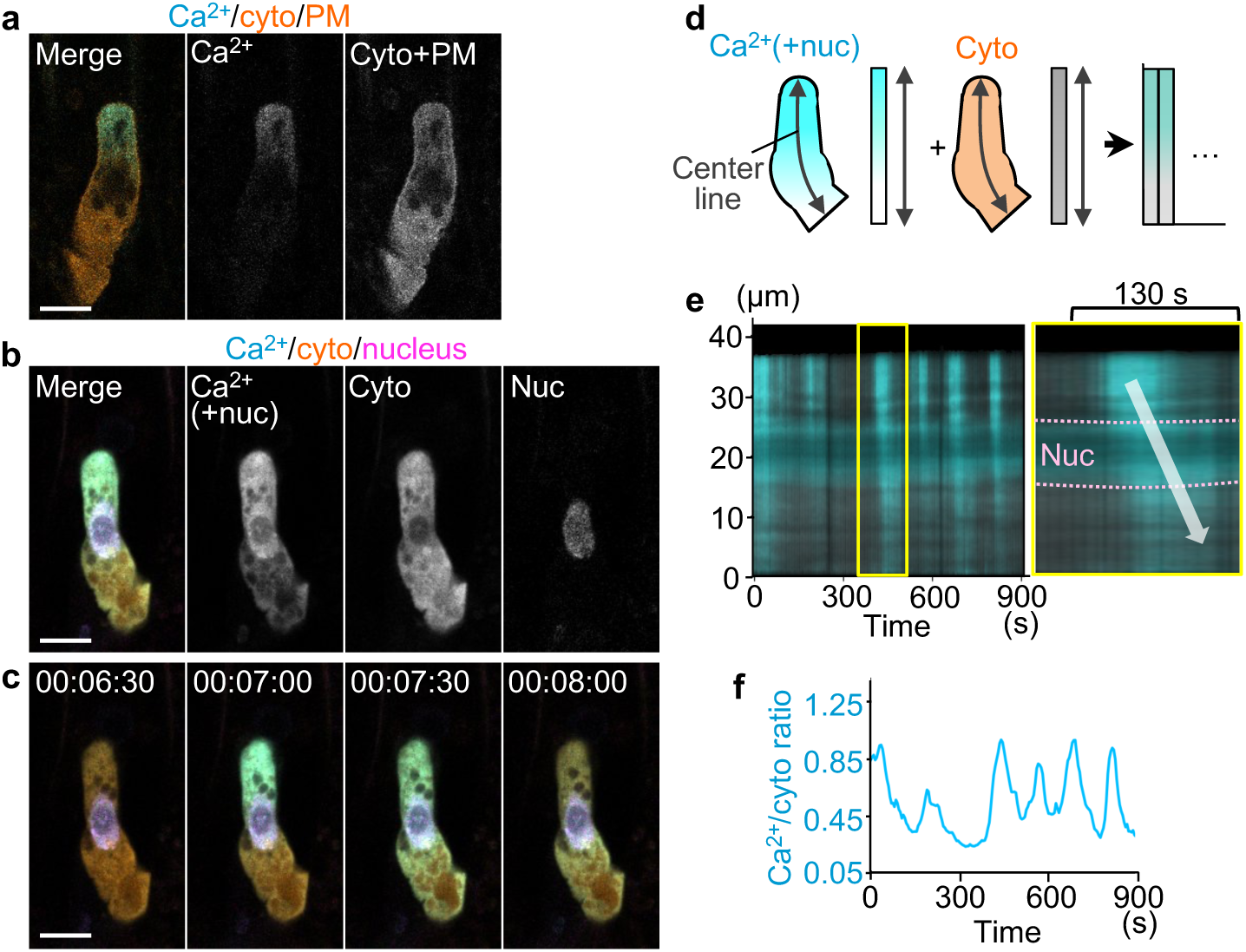
Oscillatory Ca²⁺ waves arise at the apical tip of the zygote. (**a** and **b**) Two-photon excitation microscopy (2PEM) images of the zygote expressing the Ca²⁺/cytosolic (cyto)/PM marker (**a**) or the Ca²⁺/cyto/nucleus marker (**b**). Single midplane images are shown. In **a**, the merged image (left panel) shows Ca²⁺ (cyan) and cytosolic and PM signals (cyto+PM; orange). In **b**, the merged image shows Ca²⁺ (cyan), cytosolic (orange), and nuclear (magenta) signals; the nuclear fluorescence bled into the Ca²⁺ detector channel and is therefore shown as Ca²⁺(+nuc). In both **a** and **b**, the Ca²⁺ signal appears as yellow-green due to the merge of cyan and orange. (**c**) Time-lapse 2PEM images of Ca²⁺/cyto/nucleus marker. Merged images are shown, and numbers indicate the time (h:min:s) from the first frame. (**d**) Schematics of image quantification methods used to generate a kymograph. (**e**) Kymograph showing Ca²⁺(+nuc) signal and cytosolic signal intensities projected onto the cell centerline. The right panel shows an enlarged image outlined by the yellow box. Dashed magenta lines indicate the upper and lower ends of the nucleus. White arrow indicates a single Ca²⁺ wave persisting for ∼130 s. (**f**) Graph of the Ca²⁺/cyto ratio, defined as the Ca²⁺(+nuc) signal intensity divided by the cytosolic signal intensity in whole cell. Scale bars: 10 µm.

2PEM time-lapse imaging of the Ca²⁺/cyto/nucleus marker at 5 s intervals for ≥ 10 min revealed intermittent increases and decreases in Ca²⁺ fluorescence intensity in the zygote, indicating Ca²⁺ oscillations (Fig. 1c; Supplementary Movie 1). To quantify the dynamics, we performed image analysis to extract the cell centerline and projected the fluorescence signals onto the centerline to generate kymographs (Fig. 1d; see Methods). This analysis revealed that Ca²⁺ waves initiated at the apical region of the zygote and propagated basally, and a single Ca²⁺ wave persisted for ∼130 s (Fig. 1e).

To characterize the time course, we plotted the temporal changes of Ca²⁺/cyto ratio, which is the intensity ratio of Ca²⁺(+nuc) to the cytosolic reference along the entire cell length (Fig. 1f). Although the ratio did not reach zero at any time point due to bleed-through from the nuclear marker, the oscillatory dynamics were clearly captured. The peak-to-peak interval (i.e., the oscillation period) was ∼150 s, ranging from ∼100 s (near-continuous oscillations) to ∼250 s (Fig. 1f). These periods were longer than the ∼4–50 s Ca²⁺ oscillation periods reported for pollen tubes ^13,14^.

### Ca²⁺ oscillations start with the onset of zygote elongation and attenuate before cell division

To determine whether zygotic Ca²⁺ oscillations have defined onset and offset phases during development, we imaged successive pre- and post-fertilization stages using the Ca²⁺/cyto/nucleus marker (Fig. 2). Because the Ca²⁺ oscillation period was ∼150 s, we changed the imaging intervals from 5 s to 1 min to capture Ca²⁺ peaks while avoiding photodamage from excessive laser exposure to perform long-term imaging for ≥ 10 h.

**Figure 2.**
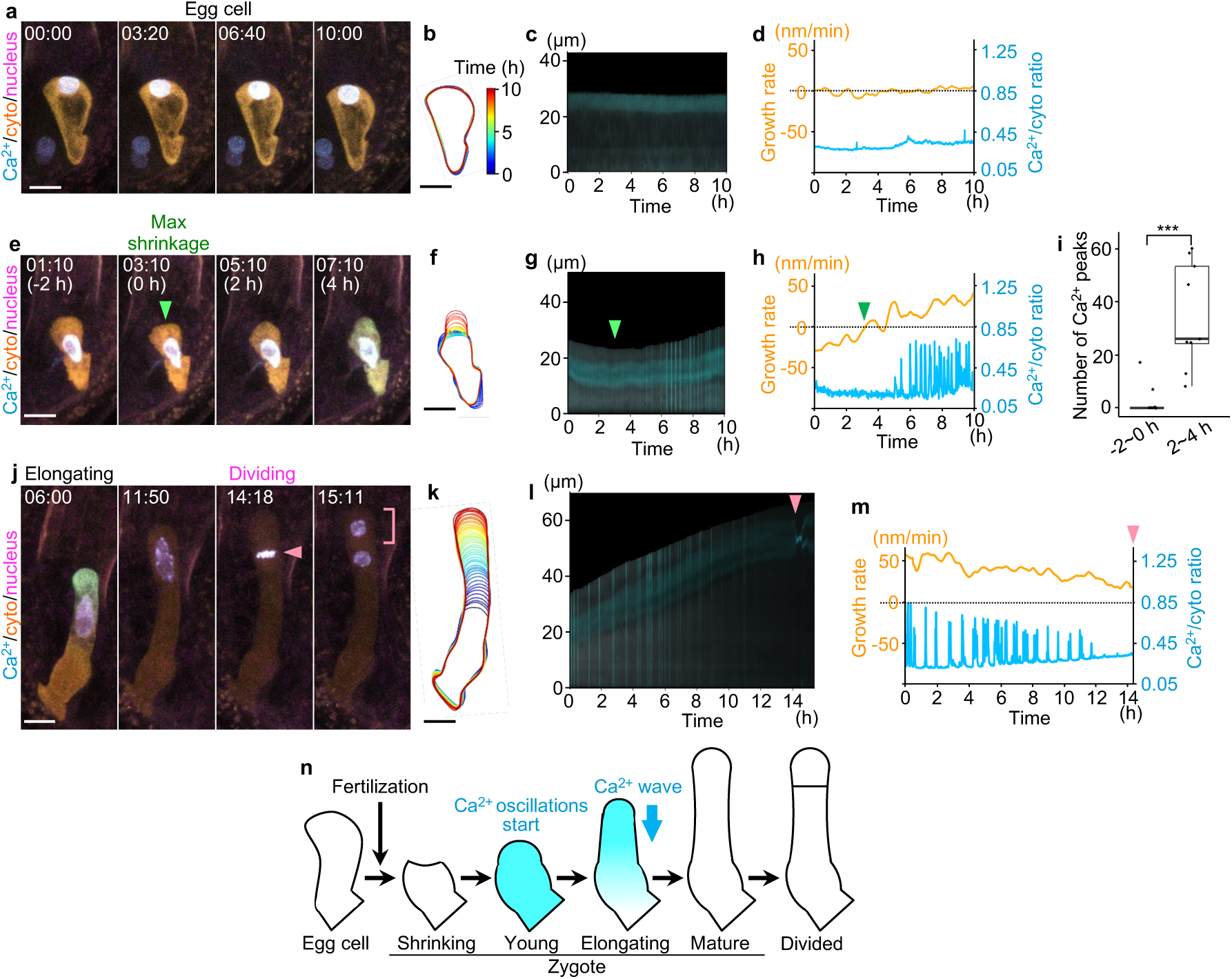
Ca²⁺ oscillations start after the onset of zygote elongation and attenuate before cell division. (**a**-**m**) Ca²⁺ oscillations and cell growth in the egg cell (**a**-**d**), young zygote (**e**-**i**), and the elongating zygote (**j**-**m**). (**a**, **e**, and **j**) Time-lapse 2PEM images of Ca²⁺/cyto/nucleus marker. Maximum intensity projection (MIP) images are shown. Numbers indicate the time (h:min) from the first frame, and the time relative to the point of maximum (max) shrinkage (defined as 0 h) is indicated in **e**. Green arrowhead in **e** indicates the cell shrinkage. Magenta arrowhead and bracket in **j** indicate the dividing nucleus and the apical cell, respectively. (**b**, **f**, and **k**) Contour dynamics of the zygote with the time indicated by color code for 10 h (**b** and **f**) and 14 h (**k**) at 20 min interval, respectively. (**c**, **g**, and **l**) Kymographs showing the signal intensity. (**d**, **h**, and **m**) Time course of cell growth rate (orange) and Ca²⁺/cyto ratio (cyan). (**i**) Boxplot of the number of Ca²⁺ peaks in 2 h before the max shrinkage (−2 to 0 h) and while after that (2 to 4 h). Significant differences were determined by the Wilcoxon rank-sum test (n = 9). \*\*\**P* ≤ 0.001. (**n**) Schematic diagram summarizing the Ca²⁺ oscillations at each stage before and after fertilization. Scale bars: 10 µm.

Before fertilization, the egg cell maintained the nucleus at the apical end, and cell shape and nuclear position did not change during time-lapse observation (Fig. 2a; Supplementary Movie 2), as shown by the unchanged cell contour (Fig. 2b) and the growth rate at ∼0 nm/min (Fig. 2d). Ca²⁺ oscillations were not detected (Fig. 2c), as shown by the Ca²⁺/cyto ratio remaining at a basal level (Fig. 2d).

We next observed young zygotes and found that the nucleus was positioned at the cell center as reported previously ^15^ (Fig. 2e; Supplementary Movie 2). The zygote first underwent transient shrinkage and then started elongation (Fig. 2f and g), as indicated by a gradual increase in growth rate from negative to positive values (Fig. 2h). Ca²⁺ oscillations began after this shrinkage phase ended and cell elongation began (Fig. 2g). The cell was too small to distinguish whether Ca²⁺ waves originated from the apical tip or the entire cell increased Ca²⁺ signal. To quantify the timing of onset, we compared Ca²⁺ oscillations before and after the time point of maximum (max) shrinkage, when the cell length was minimum (defined as 0 h; green arrowhead in Fig. 2e and g). The number of Ca²⁺ peaks during the 2 h before the max shrinkage (−2 to 0 h) was nearly zero, whereas Ca²⁺ oscillations during the 2 h after (2 to 4 h) increased significantly (Fig. 2i), indicating that Ca²⁺ oscillations start with the onset of zygote elongation.

We further imaged elongating zygotes (Fig. 2j-m; Supplementary Movie 2). Ca²⁺ oscillations persisted at high frequency during active cell growth, when the growth rate remained positive (Fig. 2l and m). Finally, elongation slowed and Ca²⁺ oscillations became rare before cell division, indicating the concurrent attenuation of Ca²⁺ oscillations and cell growth.

Together, these results indicate that Ca²⁺ oscillations begin with the onset of cell elongation and continue while elongation is active (Fig. 2n), demonstrating the close association between cell elongation and Ca²⁺ oscillations in the zygote.

### Zygotic Ca²⁺ oscillations follow cell elongation by ∼6 minutes

To test whether the temporal dynamics of Ca²⁺ oscillations and zygote growth are coupled, we performed cross-correlation analysis focusing on actively elongating zygotes (Fig. 3). To exclude bleed-through from the nuclear marker, we used the Ca²⁺/cyto/PM marker for this analysis. In addition, to capture the dynamics of Ca²⁺ oscillations with a period of ∼150 s and to simultaneously track the slow zygote growth rate (reported as < ∼70 nm/min^2^), we performed time-lapse observation at 30 s intervals for ≥ 2 h (Fig. 3a and b; Supplementary Movie 3).

**Figure 3.**
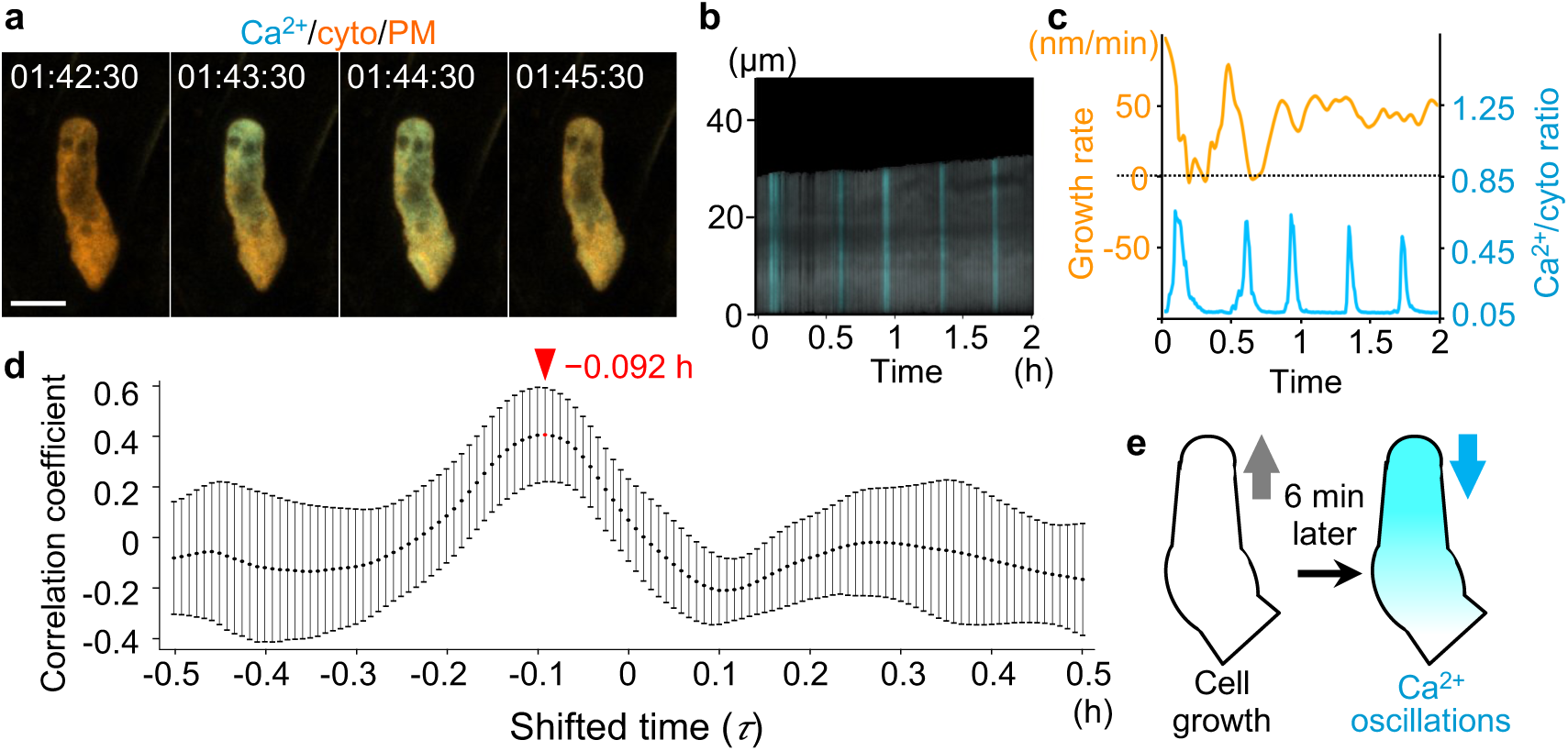
Ca²⁺ oscillations follow cell elongation by approximately 6 minutes. (**a**) Time-lapse 2PEM images of Ca²⁺/cyto/PM marker. MIP images are shown, and numbers indicate the time (h:min:s) from the first frame. (**b**) Kymograph showing the signal intensities. (**c**) Time course of cell growth rate (orange) and Ca²⁺/cyto ratio (cyan). Note that this Ca²⁺ signal does not include nuclear bleed-through; therefore, the baseline is lower than those in other figures. (**d**) Coefficient as a function of shifted time (𝜏). The red arrowhead indicates the timing (−0.092 h) with maximum. Error bars represent SD (n = 5). (**e**) Schematic diagram summarizing the time lag between cell growth and Ca²⁺ oscillations. Scale bar: 10 µm.

We focused on a 2-h period containing multiple peaks of Ca²⁺/cyto ratio (Fig. 3c), virtually shifted it relative to the fixed growth-rate trace, and calculated the correlation coefficient between the two traces as a function of the shifted time (*τ*) (Fig. 3d; Supplementary Fig. 1; see Methods). The resulting correlation coefficient was maximal at −0.092 h (∼ −6 min) (Fig. 3d), i.e., the traces aligned best when the Ca²⁺ signal was shifted ahead by ∼6 min relative to the growth-rate trace. This result indicates that Ca²⁺ oscillations recur with a consistent ∼6-min delay relative to growth-rate fluctuations during zygote elongation (Fig. 3e), suggesting that Ca²⁺ oscillations are induced by zygote growth dynamics.

### Ca²⁺ oscillations are required for zygote elongation

To investigate the role of Ca²⁺ oscillations in the zygote, we treated the Ca²⁺/cyto/nucleus marker line with the Ca²⁺ channel blocker lanthanum chloride (LaCl₃), which reduces cytosolic Ca²⁺ levels ^16^, and quantified the resulting effects (Fig. 4). To capture long-term dynamics from young to mature zygotes over ∼15 h, we performed time-lapse imaging at 20 min intervals, which could miss some Ca²⁺ peaks yet should be sufficient to capture overall Ca²⁺ oscillatory activity. Indeed, in the untreated zygote, this acquisition regime detected robust Ca²⁺ oscillations during active zygote growth, similar to those observed in Fig. 1 and 2 (Fig. 4a-c; Supplementary Movie 4).

**Figure 4.**
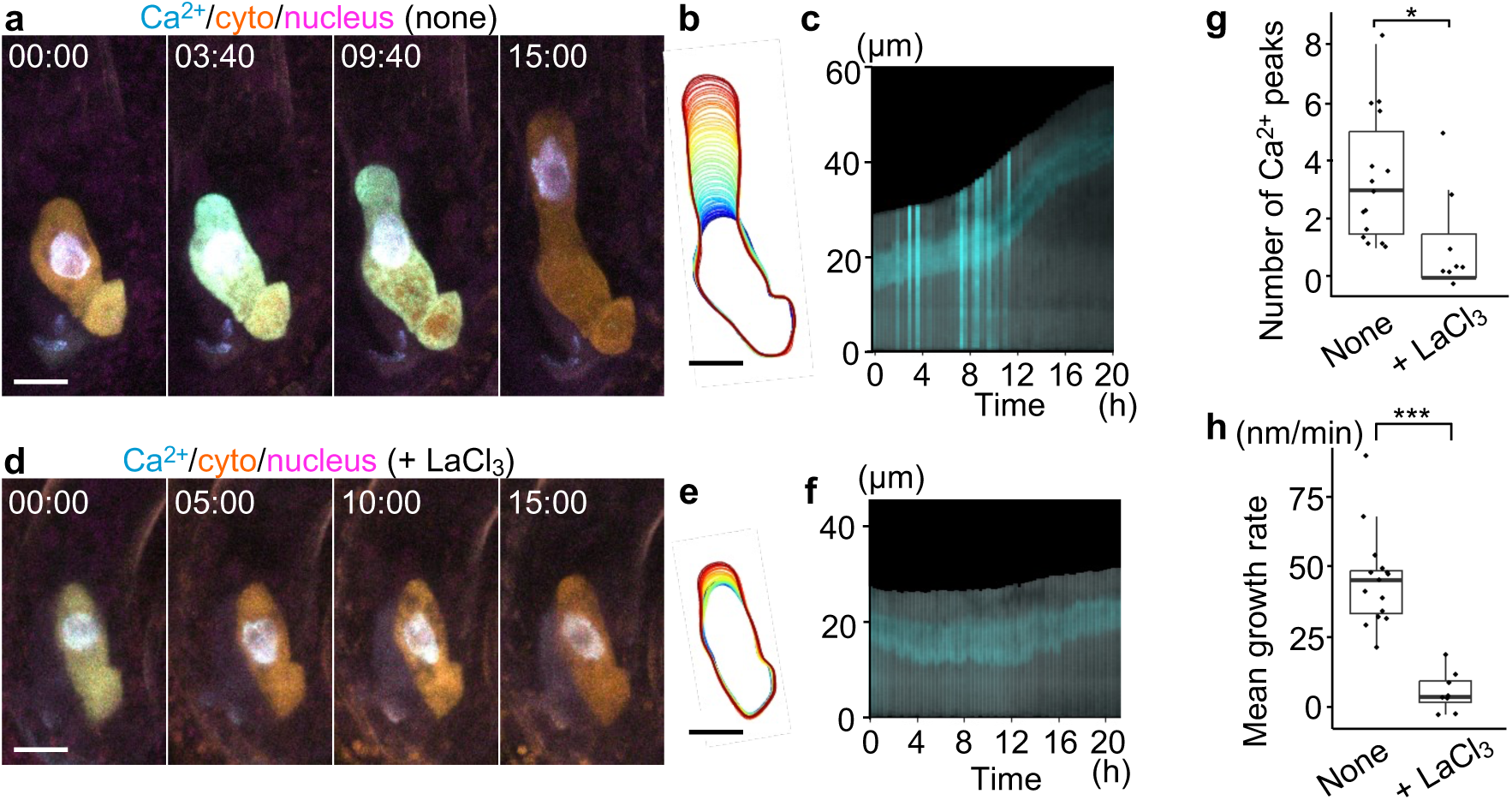
Ca²⁺ channel inhibitor blocks zygote growth. (**a**-**f**) Ca²⁺ oscillations and cell growth in untreated (none; **a**-**c**) and LaCl_3_-treated (+LaCl_3_; **d**-**f**) zygotes. (**a** and **d**) Time-lapse 2PEM images of Ca²⁺/cyto/nucleus marker. MIP images are shown, and numbers indicate the time (h:min) from the first frame. (**b** and **e**) Contour dynamics of the zygote with the time indicated by color code for 20 h at 20 min interval. (**c** and **f**) Kymographs showing the signal intensity. (**g** and **h**) Boxplots of the number of Ca²⁺ peaks (**g**) and average of growth rate (**h**) for 10 h in untreated (none) and LaCl_3_-treated zygotes. Significant differences were determined by the Wilcoxon rank-sum test (n = 15 for none and n = 8 for +LaCl_3_)). \**P* ≤ 0.05 for **g** and \*\*\**P* ≤ 0.001 for **h**. Scale bars: 10 µm.

In contrast, in the LaCl₃-treated zygote, Ca²⁺ oscillations were rarely detected and zygote elongation was severely blocked (Fig. 4d-f; Supplementary Movie 4). These effects were confirmed by the significantly reduced number of Ca²⁺ peaks (Fig. 4g) and markedly decreased mean growth rate (Fig. 4h). In addition, in approximately one-third of cases (6 of 18 zygotes), the apical tip ruptured and the cell died under LaCl₃ treatment (Supplementary Movie 4). Such growth inhibition and apical rupture are consistent with reported effects of LaCl₃ in pollen tubes, where elongation and Ca²⁺ oscillations are coupled ^17,18^.

These results indicate that Ca²⁺ oscillations are required for zygote elongation and, together with our cross-correlation analysis showing that Ca²⁺ oscillations follow zygote elongation, indicate the presence of a bidirectional feedback loop in which zygote growth and Ca²⁺ oscillations induce each other, as in other tip-growing cells ^19,20^.

### Ca²⁺ oscillations do not markedly affect overall F-actin alignment in the zygote

How do Ca²⁺ oscillations regulate zygote growth? Since Ca²⁺ signals modulate cell elongation by promoting F-actin reorganization in typical tip-growing cells ^5,6,21^, we tested whether the F-actin pattern is affected by LaCl₃ in the zygote (Supplementary Fig. 2).

We performed live imaging at 20 min intervals using an F-actin/nucleus marker line expressing an F-actin reporter (EC1p::Lifeact-mNeonGreen) and a nuclear reporter (ABI4p::H2B-tdTomato and DD22p::H2B-mCherry). In the untreated zygote, as reported previously ^7^, F-actin arrays aligned longitudinally, and this pattern was maintained during elongation (Supplementary Fig. 2a; Supplementary Movie 5). Although LaCl₃ treatment inhibited zygote elongation, F-actin filaments remained longitudinally aligned (Supplementary Fig. 2b; Supplementary Movie 5). This differs from reports in pollen tubes, where Ca²⁺ inhibition disrupts F-actin organization ^22^.

The dissociation between F-actin alignment and elongation in the zygote is consistent with previous observations that F-actin polymerization inhibitors severely impair polar organelle movements but do not cause obvious defects in zygote elongation ^7,8^. Together, these results suggest that, in the zygote, Ca²⁺ oscillations regulate cell elongation through mechanisms other than altering the overall F-actin alignment.

### Ca²⁺ oscillations prevent over-stabilization of the MT band

We then focused on the transverse MT band that supports zygotic tip growth ^9^ (Fig. 5). We performed live imaging at 20 min intervals using an MT/nucleus marker expressing an MT reporter (EC1p::Clover-TUA6) and a nuclear reporter (ABI4p::H2B-tdTomato). As reported previously ^9^, the MT band was maintained stably at the subapical region with a constant width (Fig. 5a and b; Supplementary Movie 6).

**Figure 5.**
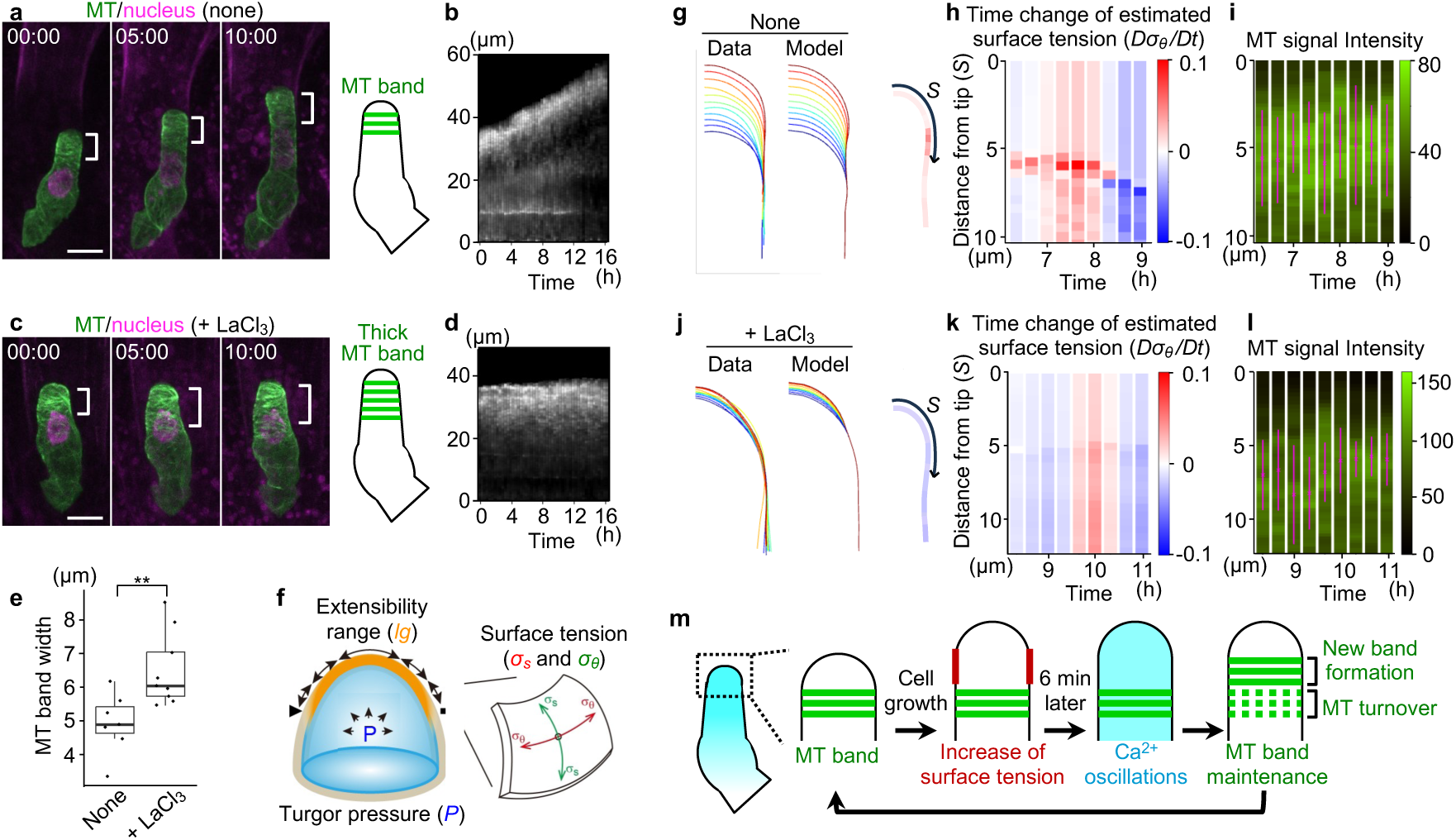
Ca²⁺ oscillations promote turnover of the microtubule (MT) band. (**a**-**d**) MT band dynamics in untreated (none; **a** and **b**) and LaCl_3_-treated (+LaCl_3_; **c** and **d**) zygotes. (**a** and **c**) Time-lapse 2PEM images of MT/nuclear marker. MIP images are shown, and numbers indicate the time (h:min) from the first frame. White brackets show the MT band, and the illustrations on the right summarize the effects. (**b** and **d**) Kymographs showing the MT signal intensity. (**e**) Boxplot of the MT band width detected by Gaussian fitting. Significant differences were determined by the Wilcoxon rank-sum test (n = 7 for none and n = 9 for +LaCl_3_). \*\**P* ≤ 0.01. (**f**) Schematic illustrations of the viscoelasto-plastic deformation model. (**g**-**l**) Results of the model analysis for the untreated zygote shown in **a** (**g**-**i**) and the LaCl_3_-treated zygote shown in **c** (**j**-**l**). (**g** and **j**) Comparison of contour dynamics of time-lapse observation data (Data) and reconstructed model zygote (Model). Only the right half of the zygote tip is shown. (**h**, **i**, **k**, and **l**) Time changes of estimated circumferential surface tension (*Dσ_θ_/Dt*) (**h** and **k**) and observed MT signal intensity (**i** and **l**). *Dσ_θ_/Dt* values at each distance from the zygote tip (*S*) are plotted with color codes. Magenta lines and crosses in **i** and **l** indicate the MT band and its center, respectively. (**m**) Schematic diagram summarizing the relation between cell growth, Ca²⁺ oscillations, and MT band organization. Scale bars: 10 µm.

In contrast, in the LaCl₃-treated zygote, the MT band became wider and broadly extended into regions farther from the tip (Fig. 5c and d). Consistently, mean values of MT band width that were detected using Gaussian fitting (see Methods) during the 10 h after imaging onset were significantly greater in LaCl₃-treated zygotes than in untreated controls (Fig. 5e). Because the MT band also thickens in zygotes treated with MT depolymerization inhibitors ^9^, these results showed the increased MT stabilization under LaCl₃ treatment and thus the effect of Ca²⁺ on destabilizing MTs.

### Ca²⁺ oscillations promote MT band turnover not via surface tension changes

Mechanical feedback at the zygote tip couples the restriction of growth direction by MT band with MT rearrangement driven by growth-dependent increase in surface tension ^9,10^. Therefore, to examine whether LaCl₃ affects this feedback, we simulated growth dynamics and mechanical changes based on the live-imaging data. Using a viscoelasto-plastic deformation model parameterized by internal expansive force (turgor pressure, *P*) and the deformable surface domain (extensibility range, *l_g_*), we calculated the spatial distribution of circumferential surface tension (one of the surface mechanical stresses in the circumferential direction: *σ_θ_*) (Fig. 5f; see Methods) ^9,23^.

We first searched for a suitable combination of *P* and *l_g_* at each time point that could reproduce the actual dynamics of zygote deformation, i.e., changes in the growth rate and tip radius obtained from the time-lapse data of the untreated zygote shown in Fig. 5a (Supplementary Fig. 3a and b), and reconstructed the model zygote (compare ‘Data’ and ‘Model’ in Fig. 5g). Based on the model zygote, we calculated the circumferential surface tension (*σ_θ_*) and its time change (*Dσ_θ_/Dt*) with respect to the surface distance from the zygote apex (*S*). Consistent with previous work ^9^, this time change markedly increased at the subapical region (Fig. 5h, red region), which corresponds to the MT band position (Fig. 5i, magenta line region).

We then derived a model using the time-lapse data from the LaCl₃-treated zygote shown in Fig. 5c (Fig. 5j; Supplementary Fig. 3c and d). Coincident with growth arrest, large time changes in the estimated surface tension were no longer detected (Fig. 5k, loss of the red region), yet the MT band remained in the subapical region (Fig. 5l, magenta line region). Thus, the MT band persisted under LaCl₃, despite growth arrest and the disappearance of estimated surface tension changes. This contrasts with the previous work, in which a mutant that ceased zygote elongation showed disappearance of the MT band within a few hours after the loss of calculated tension changes ^9^. This result suggests that Ca²⁺ oscillations promote MT band turnover via a mechanism other than surface tension changes.

## Discussion

We propose a feedback model underlying zygotic tip growth, in which oscillatory Ca²⁺ waves arise at the apical tip, tightly coupled to the cell elongation dynamics, and promote turnover of the subapical MT band that supports polar cell elongation (Fig. 5m). This model shows that the zygote employs a feedback module between Ca²⁺ oscillations and cell growth, a conserved mechanism in canonical tip-growing cells, but redirects its downstream target from the longitudinal F-actin array to the transverse MT band, thereby enabling a zygote-specific mode of tip growth.

Ca²⁺ oscillations in the zygote started when cell elongation began after the shrinkage phase, and finally attenuated before cell division (Fig. 2). In the egg cell, transient Ca²⁺ increases occur as Ca²⁺ spikes for several minutes after sperm cell discharge into the ovule and again shortly after fertilization ^24^. By contrast, the Ca²⁺ oscillations found in this work begin several hours after fertilization and persist in conjunction with elongation, suggesting that they arise from a mechanism distinct from the transient Ca²⁺ spikes. The longitudinal array of MTs in egg cells collapses during shrinkage and then reorganizes into a subapical MT band as cell elongation initiates ^7^. The temporal coincidence between elongation onset and Ca²⁺ oscillation onset may indicate that Ca²⁺-dependent regulation of the MT band starts already at the earliest phase of zygotic tip growth.

The zygote tip is actively renewed during cell elongation, and MTs accumulate in the region where growth-associated tension changes occur ^9,10^. Together with the cell growth-driven formation of a new MT band at this subapical region, Ca²⁺ oscillations may promote the clearance of older MT bands left behind it, thereby maintaining the band at a consistent position with an appropriate width (Fig. 5m). Although the molecular mechanism by which Ca²⁺ oscillations promote MT destabilization remains unknown, the zygote might utilize some Ca²⁺-dependent MT regulators. For example, in hypocotyls, Ca²⁺ elevation induces the translocation of MICROTUBULE-DESTABILIZING PROTEIN25 (MDP25) from the PM into the cytosol, which promotes MT depolymerization ^25^. Also during stomatal closure, high Ca²⁺ activates phospholipase Dα1 (PLDα1) and increases phosphatidic acid production to induce MT depolymerization ^26^. Because both MDP25 and PLDα1 are expressed in the zygote according to a public transcriptome database ^27^, these factors might connect the Ca²⁺ oscillations and MT turnover in the zygote.

Although inhibition of Ca²⁺ oscillations caused over-stabilization of the MT band even under diminished surface tension changes, surface tension may contribute to the generation of Ca²⁺ oscillations themselves. In lily pollen tubes, patch-clamp studies identified a stretch-activated Ca²⁺ current at the cell apex that responds to changes in PM extensibility, providing a model in which altered surface tension triggers Ca²⁺ influx ^28^. Therefore, zygote elongation-induced tension changes might promote the new band formation and old band clearance in a coordinated manner. In Arabidopsis pollen tubes, the mechanosensitive Ca²⁺ channel MSCS-LIKE8 has been implicated ^29^, but it is scarcely detected in the zygote transcriptome ^27^. This suggests that different Ca²⁺ channels may operate in the zygote. Indeed, the timescales of cell growth and Ca²⁺ oscillations differ markedly between pollen tubes and zygotes. Pollen tubes elongate rapidly at 2-4 µm/min, and Ca²⁺ oscillations with a ∼4-50 s period occur ∼4 s after periodic increases in growth rate ^13,14^. By contrast, the zygote elongates slowly at ≤ 70 nm/min, and Ca²⁺ oscillations with a ∼150 s period appear ∼6 min after growth fluctuations. Therefore, the Ca²⁺ channels operating in the zygote might differ in responsiveness from those in pollen tubes. Alternatively, surface-tension-independent Ca²⁺ channels might also operate in the zygote, as cyclic nucleotide-gated channels in pollen tubes can generate periodic Ca²⁺ oscillations via feedback regulation of channel activity through interactions with protein kinases and calmodulin ^30,31^. In either case, identifying the relevant Ca²⁺ channels and defining their properties in the zygote will be an important next step.

We found that Ca²⁺ oscillations in the zygote are required to regulate the transverse MT band, rather than the longitudinal F-actin array. In various well-characterized tip-growing cells, including the protonema of *Physcomitrium patens* and hyphae of fungus (*Aspergillus nidulans)*, longitudinal F-actin arrays aligned along the growth axis predominantly support sustained cell growth ^32,33^. MTs also align longitudinally in these cells, but the protonema of the fern (*Adiantum capillus-veneris*) has a transverse MT array in the subapical region, resembling the zygote MT band ^32,34,35^. To our knowledge, no study has provided clear experimental evidence that Ca²⁺ oscillations regulate transverse MTs to support tip growth. Therefore, systematic comparisons across species and cell types, e.g., between tip-growing cells with versus without an MT band, may clarify how the relationships among Ca²⁺ oscillations, the cytoskeleton, and growth dynamics are modified to support specific modes of tip growth.

## Methods

### Plant material and growth conditions

All Arabidopsis lines were in the Columbia (Col-0) background. The F-actin marker (EC1p::Lifeact-mNeonGreen; coded as MU2232) contains a green fluorescent protein mNeonGreen fused to the Lifeact sequence under the control of the *EGG CELL1* (*EC1*) promoter ^15^. The MT marker (EC1p::Clover-TUA6; coded as MU2228) contains a green fluorescent protein Clover fused with TUBULIN ALPHA6 under the control of the *EC1* promoter ^36^. The nuclear marker in the MT/nucleus marker is ABI4p::H2B-tdTomato (coded as MU2463), which expresses HISTONE H2B fused to a red fluorescent protein tdTomato under the control of the *ABA INSENSITIVE4* (*ABI4*) promoter, as described previously ^9^. The nuclear marker in the F-actin/nucleus marker contains ABI4p::H2B-tdTomato and DD22p::H2B-mCherry as zygote and endosperm nuclear reporters, respectively (coded as MU1937) ^7^. DD22p::H2B-mCherry contains DD22 promoter, H2B, and red fluorescent mCherry.

Plants were grown at 20–24°C under continuous light or long-day conditions (16-h-light/8-h-dark photoperiod).

### Plasmid construction

The Ca²⁺/cyto/nucleus marker contains EC1p::MatryoshCaMP6s as a dual-color reporter including a green fluorescent cpEGFP-based Ca²⁺ sensor (GCaMP6s) and a red/orange fluorescent LSSmOrange-based stable cytosolic marker, and EC1p::H2B-mTFP as a cyan fluorescent nuclear marker. In EC1p::MatryoshCaMP6s (coded as MU2419), the 463-bp *EC1* promoter ^37^ was fused to MatryoshCaMP6s ^12^ and NOPALINE SYNTHASE (NOS) terminator in a pMDC99 binary vector ^38^. In EC1p::H2B-mTFP (coded as MU2350), the 463-bp *EC1* promoter was fused to the coding region of HISTONE H2B (AT1G07790), a cyan fluorescent protein mTFP, and the NOS terminator in a pMDC100 binary vector38.

The Ca²⁺/cyto/PM marker contains the above-mentioned EC1p::MatryoshCaMP6s, and EC1p::tdTomato-SYP121 as a red fluorescent PM marker. In EC1p::tdTomato-SYP121 (coded as MU2490), the 463-bp *EC1* promoter was fused to a red fluorescent protein tdTomato, the coding region of SYNTAXIN RELATED PROTEIN121 (SYP121; AT3G11820), and the NOS terminator in the pPZP211 binary vector ^39^.

These constructs were transformed into Arabidopsis using the floral dip method ^40^, and at least 10 independent plants were observed in the T1 generation to assess their fluorescence pattern. At least two independent lines were established for 2PEM observation in the T2 or T3 generation.

### Time-lapse observations

The *in vitro* ovule cultivation for zygote live-cell imaging was performed using the Nitsch medium, as previously described ^41^. 2PEM images were all acquired using a laser-scanning inverted microscope (A1; Nikon) equipped with a pulse laser (Mai Tai DeepSee; Spectra-Physics). Fluorescence signals were detected by the external non-descanned GaAsP PMT detectors. Two dichroic mirrors (DM495 and DM560) were used, along with a short-pass filter (492 nm/SP) for cyan signals (mTFP), a band-pass filter (525/50 nm) for green signals (Clover, NeonGreen, and cpEGFP), and a mirror for red/orange signals (tdTomato and LSSmOrange). Time-lapse images were acquired every 5 s at 1 z-stack, every 30 s at 3 z-stacks, every 1 min at 21 or 25 stacks or every 20 min at 23 or 31 z-stacks with 1-µm intervals and using a 40× water-immersion objective lens (CFI Apo LWD WI; Nikon) with Immersol W 2010 (Zeiss) immersion medium.

For inhibitor treatment, 50 µM LaCl_3_ (Wako #127-05401) that was dissolved in water was added to the cultivation medium approximately 1 h before observation, as previously described ^42^.

Maximum intensity projection (MIP) and image processing were performed using NIS-Elements software (Nikon) or Fiji (https://fiji.sc/).

### Quantification of growth rate, Ca²⁺ signal, and MT band from time-lapse images

During live-cell imaging, ovules float in the liquid culture medium, causing fluctuations in the zygote position. We therefore performed coordinate normalization and apical tip tracking, as previously described ^2,43^. This processing enabled extraction of the cell centerline from the basal end to the apical tip, and the time change in cell length was calculated as growth rate.

For Ca²⁺ quantification, the fluorescence intensity of the above-mentioned green signals was defined as [Ca²⁺] (Ca²⁺(+nuc) for the Ca²⁺/cyto/nucleus marker, and Ca²⁺ for the Ca²⁺/cyto/PM marker), and the fluorescence intensity of the above-mentioned red signals was defined as [Cyt]. For both signals, intensity values were averaged over the whole-cell area, i.e., interior region within the segmented cell contour throughout the study. The Ca²⁺/cyto ratio was then calculated as [Ca²⁺]/[Cyt]. For the number of Ca²⁺ peaks, we counted peaks in the Ca²⁺/cyto ratio that exceeded a threshold defined as the grand mean of the mid-range values (the average of the maximum and minimum) calculated for each sample.

For MT quantification, MT signal intensity within the segmented cell contour was projected onto the cell centerline. The resulting one-dimensional intensity profile was fitted with a truncated Gaussian function to maximize the coefficient of determination. We then estimated the MT band position and width corresponding to the mean and the double of standard deviation, as previously described ^23^.

### Cross-correlation analysis of growth rate and Ca²⁺/cyto ratio

The analysis workflow is summarized in Supplementary Fig. 1. We used data of growth rate and Ca²⁺/cyto ratio containing at least 4 peaks within a 2 h window (Supplementary Fig. 1a). To remove high frequency noise in the growth rate, it was smoothed using locally weighted scatterplot smoothing (LOWESS) with time window of 900 s, which is relatively higher than the oscillatory period (∼150 s) of the Ca²⁺/cyto ratio. To quantify correlation, we subtracted the mean and normalized the series by the standard deviation, and the resulting series were defined as the adjusted 𝑋(𝑡) (growth rate) and 𝑌(𝑡) (Ca²⁺/cyto ratio) (Supplementary Fig. 1b). We fixed 𝑋(𝑡) and generated shifted series 𝑌(𝑡 + 𝜏) by exhaustively shifting 𝑌(𝑡) from 𝜏 = −0.5 to +0.5 h (Supplementary Fig. 1c). We applied the cross-correlation coefficient 𝜌*_XY_*(𝜏) to these time series as follows (Supplementary Fig. 1d).

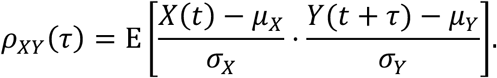

where 𝜇*_X_* and 𝜇*_y_* denote the time averages of 𝑋(𝑡) and 𝑌(𝑡), respectively, and 𝜎*_X_* and 𝜎*_Y_* denote the standard deviations of 𝑋(𝑡) and 𝑌(𝑡), respectively. The operator 𝐸[⋅] stands for the mean of the variable ⋅. We performed the same analysis for all samples to determine the 𝜏 that maximized the correlation coefficient.

### Estimation of surface tension based on viscoelasto-plastic deformation model

We built a mechanical model for zygote elongation based on the model previously developed ^9,23,44^. This model characterizes the shape, stress and strain to calculate the next cell shape. Cell surface mechanical stresses in the meridional and circumferential directions were denoted by 𝜎*_S_* and 𝜎_θ_ and calculated with the input of turgor pressure 𝑃, the cell wall thickness, and the meridional and circumferential curvature (see details in Kang et al. 2024 ^23^). The strain rates on the surface in the meridional and circumferential directions were then calculated by the cell surface extensibility and some mechanical parameters ^44^. The extensibility is determined by the extensible range 𝑙*_g_* and the magnitude of extensibility at the tip (same notation in Kang et al. 2024 ^23^). The displacement vectors in the tangential and perpendicular direction were finally estimated. The initial configuration is based on the hemispherical shape with the radius larger than that in the data. The simulation step was smaller than the data time interval and we solved the above equations per simulation step. We estimated the parameter set (𝑃, 𝑙*_g_*) through the quantification of growth rate and tip radius 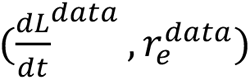 (ellipse fitting in Kang *et al. 2024 ^9^). We exhaustively searched for the best fitted model parameters (𝑃, 𝑙*_g_*) fitted with 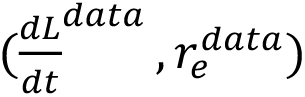 and resulting 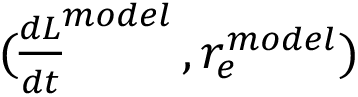 that minimizes the error function 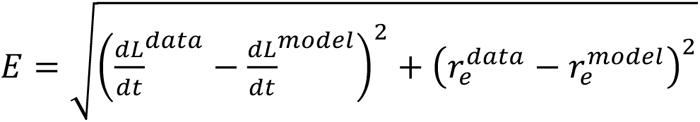. This fitting process was applied to each pair of sequential time frames.

The time change of the circumferential stress was calculated as,

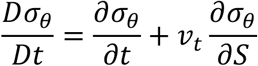

where 𝑣 is the displacement vector in the tangential direction. We note that the time partial derivative is corrected with the tangential displacement depending on the Lagrangian flow of the point on the cell surface.

### Quantification and statistical analysis

Statistical analysis was performed using R and Python. The methods used for the statistics and the number of samples are described in the respective legend.

## Data availability

Data supporting the findings of this work are available within the paper and its Supplementary files. Plasmids and transgenic lines generated in this study are available upon reasonable request to the corresponding author.

## Code availability

No new code was generated in this paper. All analytical methods are described in the Methods section.

## Acknowledgements

We thank Hisa Yoshida, Tamiko Ambo, Satomi Watanabe, Yuko Kudo and Junko Kato for technical support, Tetsuya Higashiyama for helpful discussions, and Wolf B. Frommer for providing MatryoshCaMP marker. This work was supported by the Japan Society for the Promotion of Science [a Grant-in-Aid for Early-Career Scientists (JP22K15135 and JP25K18484 to H.M., JP23K14204 to Y.K., and JP25K18499 to Z.K.), Grant-in-Aid for Transformative Research Areas (A) (JP25H01809 to Y.K.), a Grant-in-Aid for Scientific Research on Innovative Areas (JP16H06280 (Advanced Bioimaging Support)), a Grant-in-Aid for Scientific Research (B) (JP23H01143 to S.T. and JP23H02494 to M.U.), and International Leading Research (JP22K21352) to M.U.], the Japan Science and Technology Agency [CREST (JPMJCR2121 to S.T. and M.U., YORC to H.M., Y.K., Z.K. and T.N.)], the Inamori Foundation (Inamori Research Grant to Y.K.), and the Suntory Rising Stars Encouragement Program in Life Sciences (SunRiSE to M.U.).

## Author Contributions

H.M. and M.U. designed the research; H.M. carried out the experiments; H.M., Z.K., and T.N. analyzed the data; Z.K., T.N., Y.K., and S.T. contributed materials/analysis tools; and H.M., S.T., and M.U. wrote the manuscript.

