## Supplementary Fig for "Ca²⁺ oscillations promote microtubule-band turnover and support tip growth in Arabidopsis zygotes"

**Supplementary materials include:**
Supplementary Figures 1 to 3
Supplementary Movies 1 to 6

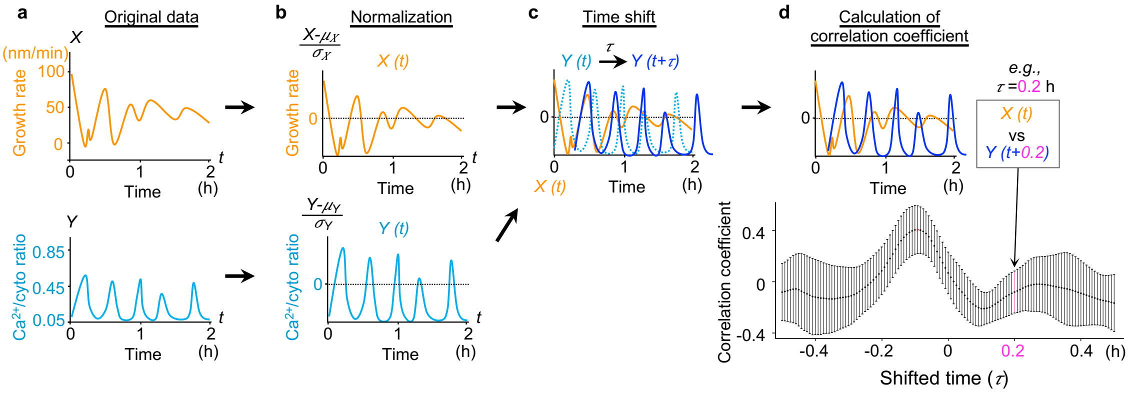


**Supplementary Figure 1. Procedure of cross-correlation analysis.**

Analysis workflow to obtain cross-correlation coefficient of cell growth and $Ca^{2+}$ oscillations. (**a**) Original data of growth rate (orange; $X$) and the $Ca^{2+}$/cyto ratio (cyan; $Y$). (**b**) Normalization of growth rate ($X(t)$) and $Ca^{2+}$/cyto ratio ($Y(t)$) by subtracting mean and dividing by standard deviation. (**c**) Time shift of $Y(t)$ to $Y(t+\tau)$ by $+\tau$ against the fixed $X(t)$. (**d**) Calculation of the correlation coefficient between $X(t)$ and $Y(t+\tau)$ as a function of shifted time ($\tau$).

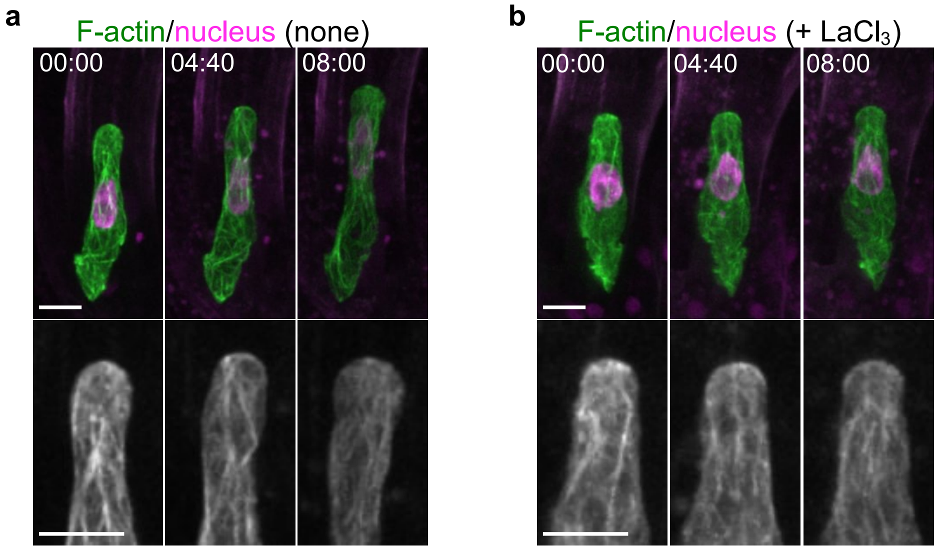


**Supplementary Figure 2. Effects of $Ca^{2+}$ channel inhibitor on F-actin dynamics.**

Time-lapse 2PEM images of F-actin/nucleus marker in untreated (none; **a**) and $LaCl_3$-treated (+$LaCl_3$; **b**) zygotes. Upper and lower panels show merged images and enlarged F-actin images, respectively. MIP images are shown, and numbers indicate the time (h:min) from the first frame. Scale bars: 10 μm.

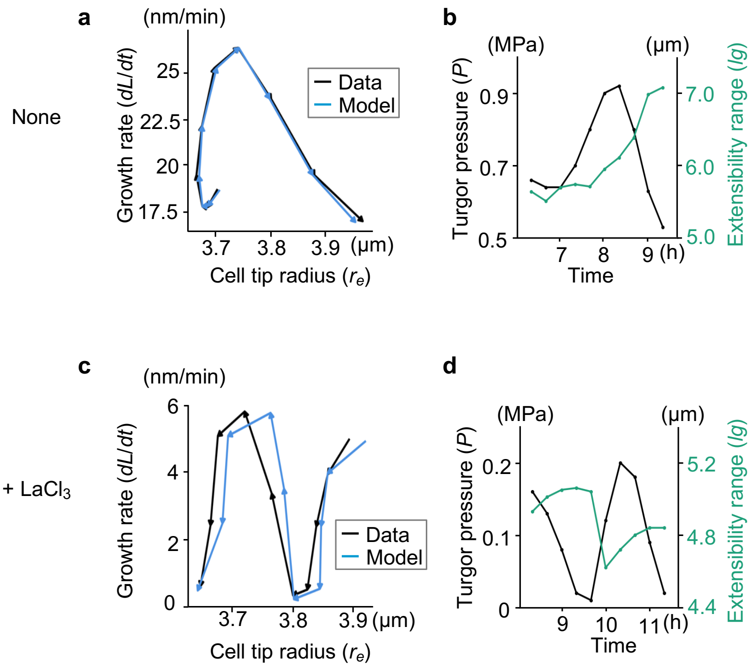


**Supplementary Figure 3. Parameter estimation using viscoelasto-plastic deformation model.**

(**a**-**d**) Model analysis for the untreated zygote shown in Fig. 5**a** (none; **a** and **b**) and the $LaCl_3$-treated zygote shown in Fig. 5**c** (+$LaCl_3$; **c** and **d**). (**a** and **c**) Comparisons of growth rate (*dL/dt*) and cell tip radius ($r_e$) between time-lapse observation data (Data; black) and reconstructed model zygote (Model; blue). (**b** and **d**) Estimated turgor pressure (*P*; black) and extensibility range ($l_g$; green).

**Legend for Supplementary Movies**

**Supplementary Movie 1. Dynamics of $Ca^{2+}$ wave in the elongating zygote.**

Two-photon excitation microscopy (2PEM) time-lapse images of the zygote expressing the $Ca^{2+}$ (cyan), cytosolic (orange), and nuclear (magenta) marker. The images were acquired at 5 s intervals, and numbers indicate the time (h:min:s) from the first frame. Single midplane images are shown. Images are representative of 4 time-lapse images. Scale bar: 10 μm.

**Supplementary Movie 2. Dynamics of $Ca^{2+}$ oscillations at each stage before and after fertilization.**

2PEM time-lapse images of a zygote expressing the $Ca^{2+}$ (cyan), cytosolic (orange), and nuclear (magenta) marker in the egg cell (left), young zygote (middle), and the elongating zygote (right). The images were acquired at 1 min intervals, and numbers indicate the time (h:min) from the first frame. MIP images are shown. Images are representative of 5, 9, and 7 time-lapse images for the egg cell, young zygote, and elongating zygote, respectively. Scale bar: 10 μm.

**Supplementary Movie 3. Dynamics of $Ca^{2+}$ oscillations in the elongating zygote.**

2PEM time-lapse images of a zygote expressing the $Ca^{2+}$ (cyan), cytosolic (orange), and PM (orange) marker in the elongating zygote. The images were acquired at 30 s intervals, and numbers indicate the time (h:min:s) from the first frame. MIP images are shown. Images are representative of 5 time-lapse images. Scale bar: 10 μm.

**Supplementary Movie 4. Effects of $LaCl_3$ on $Ca^{2+}$ oscillations and cell dynamics.**

2PEM time-lapse images of a zygote expressing the $Ca^{2+}$ (cyan), cytosolic (orange), and nuclear (magenta) marker in untreated (none; left) and $LaCl_3$-treated (+$LaCl_3$; middle and right) zygotes. Middle and right movies show arrested and ruptured zygotes, respectively. The images were acquired at 20 min intervals, and numbers indicate the time (h:min) from the first frame. MIP images are shown. Images are representative of 15, 8, and 6 time-lapse images for untreated, and arrested and ruptured zygotes among the $LaCl_3$-treated zygotes, respectively. Scale bar: 10 μm.

**Supplementary Movie 5. Effects of $LaCl_3$ on F-actin dynamics.**

2PEM time-lapse images of a zygote expressing the F-actin (green) and nuclear (magenta) marker in untreated (none; left) and $LaCl_3$-treated (+$LaCl_3$; right) zygotes. The images were acquired at 20 min intervals, and numbers indicate the time (h:min) from the first frame. MIP images are shown. Images are representative of 5 and 5 time-lapse images for untreated and $LaCl_3$-treated zygotes, respectively. Scale bar: 10 μm.

**Supplementary Movie 6. Effects of $LaCl_3$ on MT dynamics.**

2PEM time-lapse images of a zygote expressing the microtubule (MT; green) and nuclear (magenta) marker in untreated (none; left) and $LaCl_3$-treated (+$LaCl_3$; right) zygotes. The images were acquired at 20 min intervals, and numbers indicate the time (h:min) from the first frame. MIP images are shown. Images are representative of 7 and 9 time-lapse images for untreated and $LaCl_3$-treated zygotes, respectively. Scale bar: 10 μm.
